# Redistribution of the novel *C. difficile* spore-adherence receptor, E-cadherin, by TcdA and TcdB increases spore-binding to adherens junctions

**DOI:** 10.1101/2021.09.01.458577

**Authors:** Pablo Castro-Córdova, Macarena Otto-Medina, Nicolás Montes-Bravo, Christian Brito-Silva, D. Borden Lacy, Daniel Paredes-Sabja

**Affiliations:** ANID – Millennium Science Initiative Program – Millennium Nucleus in the Biology of the Intestinal Microbiota, Santiago, Chile; Departamento de Ciencias Biológicas, Facultad de Ciencias de la Vida, Universidad Andrés Bello, Santiago, Chile; Department of Pathology, Microbiology, and Immunology, Vanderbilt University School of Medicine, Nashville, Tennessee, USA; Department of Biology, Texas A&M University, College Station, Texas, USA

**Keywords:** *C. difficile* spores, E-cadherin, adherens junctions, spore-host interaction, *C. difficile* toxins TcdA/TcdB

## Abstract

*Clostridioides difficile* causes antibiotic-associated diseases in humans ranging from mild diarrhea to severe pseudomembranous colitis and death. A major clinical challenge is the prevention of disease recurrence, which affects nearly ∼20 to 30 % of the patients with a primary *C. difficile* infection (CDI). During CDI, *C. difficile* forms metabolically dormant spores that are essential for recurrence of CDI (R-CDI). In prior studies, we have shown that *C. difficile* spores interact with intestinal epithelial cells (IECs), which contributes to R-CDI. However, this interaction remains poorly understood. Here, we provide evidence that *C. difficile* spores interact with E-cadherin, contributing to spore-adherence and internalization into IECs. *C. difficile* toxins TcdA/TcdB lead to adherens junctions opening and increase spore-adherence to IECs. Confocal micrographs demonstrate that *C. difficile* spores associate with accessible E-cadherin; spore-E-cadherin association increases upon TcdA/TcdB intoxication. The presence of anti-E-cadherin antibodies decreased spore adherence and entry into IECs. By ELISA, immunofluorescence, and immunogold labelling, we observed that E-cadherin binds to *C. difficile* spores, specifically to the hair-like projections of the spore, reducing spore-adherence to IECs. Overall, these results expand our knowledge of how *C. difficile* spores bind to IECs by providing evidence that E-cadherin acts as a spore-adherence receptor to IECs and by revealing how toxin-mediated damage affects spore interactions with IECs.

## INTRODUCTION

*C. difficile* is a Gram-positive, anaerobic, spore-forming bacterium and the main causative agent of antibiotic-associated diarrhea, causing death in ∼5% of cases (1, 2). R-CDI is a major clinical challenge affecting ∼30% of CDI patients (2, 3). CDI is primarily driven by the *C. difficile* toxins TcdA and TcdB, with contributions from the binary transferase toxin in some cases (4-6). Both TcdA and TcdB target their intracellular glucosyltransferase activity towards RhoA, Rac1, and Cdc42 (7, 8). This inhibitory modification of the Rho GTPase family leads to actin depolymerization and disruption of tight and adherens junctions (9-13), causing disruption of epithelial paracellular barrier function (14). Clinical studies showed that administration of anti-TcdB antibody (bezlotoxumab) in combination with antibiotics reduces R-CDI rates (15). The antibody may prevent toxin-mediated stem cell dysfunction and therefore prevent the delay in recovery and repair of the intestinal epithelium (10).

During infection, *C. difficile* produces spores (16). Strains unable to form spores do not cause R-CDI in animal models (17), indicating that the spores are critical for R-CDI. EDTA-disruption of cell-cell junctions in Caco-2 cells leads to increased adherence of *C. difficile* spores at the cell-cell junction, suggesting a tropism for cell-cell junctions (18). *C. difficile* spores bind to fibronectin and vitronectin, located in the basolateral membrane (19). These molecules contribute to *C. difficile* spore internalization into IECs in an integrin-dependent manner, and spore-entry *in vivo* contributes to the R-CDI (20).

The intestinal barrier is an impermeable layer that separates the apical and basolateral cell sides (21), and is formed by junctions between adjacent IECs. The cell-cell junctions between IECs are constituted of E-cadherin containing adherens junctions and tight junctions formed by claudins, occludins, and zonula occludens. These junctions provide the gut barrier with impermeability properties and restrict the translocation of pathogens and commensal bacteria. However, some pathogens can alter or disrupt the cell junctions, to translocate through the host (22).

Several enteric pathogens such as *Streptococcus pneumoniae, Fusobacterium nucleatum, Campylobacter jejuni*, and *Listeria monocytogenes* take advantage of disrupted intestinal adherens junctions to internalize in host cells (23-27). How *C. difficile* spores interact with adherens junctions remains poorly described. Herein, we show that accessible E-cadherin is reduced in differentiated Caco-2 cells. Also, TcdA/TcdB intoxication increases the spore adherence that is correlated with an increase in E-cadherin accessibility in IECs. Next, we observed that *C. difficile* spores interact with E-cadherin in healthy IECs and the binding is increased in TcdA/TcdB-intoxicated cells. Also, the blocking of the cellular E-cadherin with monoclonal antibodies reduces the spore adherence and entry to IECs. Finally, we observed that E-cadherin binds to the hair-like extensions of the exosporium.

## METHODS

### Bacterial growth conditions

*C. difficile* was grown under anaerobic conditions at 37 °C in a Bactron III–2 chamber (Shellab, OR, USA) in BHIS medium; 3.7% brain–heart infusion broth (BD, USA) supplemented with 0.5% of yeast extract (BD, USA), and 0.1% L–cysteine (Merk, USA) liquid or on plates with 1.5% agar (BD, USA). *C. difficile* strain R20291 wild-type B1/NAP1/PCR-ribotype 027 strain obtained from CRG0825 stock at Nottingham Clostridia Research Group (28). *C. difficile* 630Δ*erm* is a toxigenic (*tcdA*^+^ *tcdB*^+^) strain of *C. difficile* isolates from a patient with pseudomembranous colitis during an outbreak of CDI (29). *C. difficile* 630Δ*erm bclA1::CT*, 630Δ*erm bclA2::CT* and 630Δ*erm bclA3::CT* are insertional inactivation clostron mutant strains in the exosporium collagen-like proteins BclA1, BclA2 and BclA3 (30), and were a kind gift from Dr. Simon Cutting (School of Biological Sciences, Royal Holloway, University of London). *Bacillus megaterium* transformed with the plasmid pHIS 1622 that contains the cloned genes *tcdA* and *tcdB* of *C. difficile* VPI 10463 (31), were aerobically grown at 37 °C in Luria-Bertani (BD) with 10 µg/mL tetracycline.

### Cell lines and culture conditions

Caco-2 cells (ATCC, USA) were grown at 37 °C with 5% CO_2_ with Dulbecco’s modified Eagle’s minimal essential medium High Glucose (DMEM; Hyclone, USA) supplemented with 10% of fetal bovine serum (FBS; Hyclone, USA) and 50 U mL^−1^ penicillin, and 50 µg mL^−1^ streptomycin (Corning Cellgro, USA) as previously has been described For experiments, cells were plated over glass coverslips in 24–well plates and were cultured for 2–days post–confluence (undifferentiated) or 8–days post–confluence (differentiated), changing the media every other day. To polarize Caco-2 cells were cultured onto 12mm Transwell (Corning, USA) with pores size of 0.4 µm until transepithelial electric resistance >250 Ω × cm^2^.

### Recombinant *C. difficile* toxin Purification

*C. difficile* toxins, TcdA and TcdB, were expressed in *B. megaterium* transformed with the plasmid pHIS1622, containing the cloned genes *tcdA* and *tcdB* of *C. difficile* VPI 10463 as previously described (31). Briefly, transformed *B. megaterium* were cultured in Luria–Bertani medium (BD, USA) supplemented with 10 µg mL^−1^ tetracycline, and 35 mL of an overnight culture was used to inoculate 1L of media. Bacteria were grown at 37 °C with 220 rpm. Toxin expression was induced with 0.5% D–xylose once the culture reached OD_600_ = 0.5. Cells were harvested after 4 h by centrifugation at 4,750×*g* and resuspended in 200 mL of binding buffer (20 mM Tris [pH 8.0], 100 mM NaCl for TcdA and 20 mM Tris [pH 8.0], 500 mM NaCl for TcdB) supplemented with 0.16 µg mL^−1^ DNase, 10 mg mL^−1^ lysozyme and protease inhibitors (P8849; Sigma). Cell lysis was performed using an Emulsiflex homogenizer, and lysates were centrifuged at 48,000×*g* for 30 min. The toxins were purified from the supernatant by Nickel–affinity, anion exchange, and size–exclusion chromatography. Toxins were eluted and stored in 20 mM HEPES (pH 7.0), 50 mM NaCl.

### Spore preparation and purification

*C. difficile* spores were purified as we previously described (32). Briefly, *C. difficile* spore was cultured overnight in BHI medium in a Bactron III–2 anaerobic chamber (Shellab, USA). Then a 1:500 dilution was plated in TY agar plates; 3% trypticase soy and 0.5% yeast extract and 1.5% Bacto agar (BD, USA) and incubated for 7 days at 37 °C in anaerobiosis. Colonies were harvested with sterile, cold Milli–Q water, washed 3 times and purified with 50% Nicodenz. The spore suspension was 99% free of vegetative cells, sporulating cells, and debris. Spores were quantified in a Neubauer chamber and stored at −80 °C until use.

### Immunofluorescence of cell monolayers

To evaluate E-cadherin distribution in undifferentiated and differentiated Caco-2 cells, cells were washed with PBS and were fixed with PBS–4% paraformaldehyde for 20 min at 4 °C, blocked with PBS–3% BSA 1 h at room temperature (RT). To stain the accessible E-cadherin, cells were incubated with 1: 50 of primary antibody mouse polyclonal anti–human E-cadherin (sc-8426, Santa Cruz Biotechnology, USA) in PBS–3% BSA, for 2 h at RT followed by two washed with PBS and 1:200 secondary antibody donkey anti–mouse-Alexa Fluor 568 (ab175700, Abcam, USA). To detect the total E-cadherin, fixed samples were permeabilized with PBS-0.2% Triton 100-X for 10 min at RT and blocked with PBS–3% BSA for 1 h at RT. The total protein was detected with the same primary antibody used accessible E-cadherin; primary antibody mouse polyclonal anti–human E-cadherin (sc-8426, Santa Cruz Biotechnology, USA) in PBS–3% BSA that was incubated for 2 h at RT, then were washed and was incubated with 1:200 secondary antibody donkey anti–mouse-Alexa Fluor 568 (ab175700, Abcam, USA). The non–permeabilized samples stained for accessible E-cadherin were permeabilized with PBS-0.2% Triton 100-X for 10 min at RT, washed, and to stain the F–actin, cells were incubated with 1:1,000 Phalloidin i– Fluor 488 (ab176753, Abcam, USA) in PBS for 30 min at RT. Cells were washed, and nuclei were stained with 8 µg mL^−1^ Hoechst 33342 (ThermoFisher, USA) in PBS for 10 min at RT. Then samples were washed and mounted using Dako Fluorescent Mounting Medium (Dako, Denmark). Samples were visualized in the confocal microscope Leica TCS LSI (Leica, Germany). Please see below.

To study the redistribution of E-cadherin in TcdA/TcdB intoxicated cells, differentiated Caco-2 cells cultured into 24–well plates were washed twice with DPBS (Hyclone, USA) and were intoxicated with 600 pM TcdA and 600 pM TcdB for 3, 6, and 8 h in DMEM, without serum. As a control, cells were incubated in the same solution without TcdA/TcdB. Then cells were rinsed with PBS, fixed with PBS–4% paraformaldehyde, and washed with PBS and blocked with PBS– 1% BSA at 4 °C overnight. To stain the surface, accessible E-cadherin cells were incubated with 1:200 mouse mAb anti–E-cadherin (SC8426, Santa Cruz Biotechnologies, USA), 1 h at RT, washed, and followed incubation with 1:400 donkey anti–mouse IgG Alexa Fluor 488 (ab150109, Abcam, USA) for 1 h at RT. To detect the total protein, samples were permeabilized with PBS– 0.2% Triton 100-X for 10 min at RT and blocked with PBS–1% BSA for 1 h at RT. The total protein was detected with the same primary antibody used before, washed, and samples were incubated with donkey anti–mouse IgG Alexa Fluor 568 (ab175700, Abcam, USA). Samples were mounted using Dako Fluorescent Mounting Medium (Dako, Denmark) and were visualized in the confocal microscope Leica Sp8 (Leica, Germany; please see below.)

To evaluate *C. difficile* spore association with accessible E-cadherin in TcdA/TcdB intoxicated IECs, differentiated Caco-2 cells were intoxicated with 600 pM TcdA and 600 pM TcdB for 8 h at RT, as described above. Then cells were infected with *C. difficile* spores in an MOI of 10 for 1 h at 37 °C. Then cells were washed, fixed with PBS–4% paraformaldehyde for 10 min at RT, blocked with PBS–1% BSA overnight at 4 °C, and *C. difficile* spores were stained with 1:1,000 primary polyclonal anti–*C. difficile* spore IgY batch 7246 antibodies (33) (Aveslab, USA) in PBS–1% BSA for 1 h at RT. Following two PBS washed, samples were incubated with 1:400 goat anti–chicken IgY secondary antibodies Alexa–Fluor 488 (#ab150173 Abcam, USA) in PBS– 3% BSA for 1 h at RT, washed two times with PBS, and accessible E–cadherin was stained as it is described above. Cellular nuclei were stained with 8 µg mL^−1^ Hoechst 33342 (ThermoFisher, USA) for 15 min at RT. Finally, samples were mounted and analyzed by confocal microscopy Leica SP8.

The Leica microscopes TCS LSI and SP8 at the Confocal Microscopy Core Facility of the Universidad Andrés Bello were used. To evaluate E-cadherin distribution in undifferentiated and differentiated Caco-2 cells, Leica TCS LSI was used with 63× ACS APO oil objective numerical aperture 1.3, with an optical zoom of 2×. Micrographs were acquired using excitation wavelengths of 405, 488, and 532 nm, and signals were detected with an ultra–high dynamic photomultiplier (PMT) spectral detector (430–750nm). Emitted fluorescence was split with four dichroic mirrors (QD 405, 488, 561, and 635 nm). Images 1,024 × 1,024 pixels were acquired each 0.3 µm. To observe the redistribution of accessible E-cadherin and total E-cadherin in TcdA and TcdB intoxicated cells, Leica SP8 was used with HPL APO CS2 40× oil, numerical aperture 1.30. For signal detection, three PMT spectral detectors were used; PMT1 (410–483) DAPI, PMT2 (505–550), Alexa–Fluor 488, and PMT3 (587–726) Alexa–Fluor 555. Emitted fluorescence was split with dichroic mirrors DD488/552. Pictures of 1,024 × 1,024 pixels were captured each 0.3 µm. Images of 3–dimensional reconstructions of the Caco-2 monolayers were performed using ImageJ software (NIH, USA). In undifferentiated and differentiated cells, accessible E-cadherin cells were quantified as the number of cells with positives staining. To quantify the redistribution of E-cadherin, Fl. Int. of each slide was measured using the plug–in “Measure stack” of Fiji (ImageJ, NHI, USA), and the sum of the Fl. Int. of each slide is considered as the Fl. Int. of the *z*–stack of a field for accessible and total E-cadherin. For each field, the same number of slides was used.

### Quantification of *C. difficile* spore adherence and internalization

To evaluate spore adherence and internalization in TcdA/TcdB intoxicated cells, differentiated Caco-2 cells were incubated with 600 pM TcdA and 600 pM TcdB for 3, 6, and 8 h in DMEM, without serum. As a control, cells were incubated in the same solution without TcdA/TcdB or cells were incubated with TcdA/TcdB previously heat inactivated at 70°C for 5 min. Then cells were infected with *C. difficile* spores an MOI 10 for 3 h at 37 °C. Unbound spores were rinsed off with PBS, and cells were fixed with PBS–4% paraformaldehyde for 10 min, and adhered spores were stained as described below.

To evaluate spore adherence and entry in presence of anti–E-cadherin, undifferentiated, differentiated, and polarized Caco-2 cells were incubated with 0,1; 1,0; 10; 100 µg mL^−1^ of mouse monoclonal anti–human E-cadherin (SC8426, Santa Cruz Biotechnology, USA) or as control non– immune IgG antibody of rabbit serum (I5006, Sigma-Aldrich, USA) for 2 h at 37 °C. Then cells were infected for 3 h at 37 °C with *C. difficile* spores, previously incubated with 20 µL of FBS per well for 1 h at 37 °C. Cells were washed two times with PBS to remove unbound spores and were fixed with PBS–4% paraformaldehyde for 10 min at RT, and adhered spores were stained as described below.

To evaluate spore adherence and entry in presence of purified E-cadherin, undifferentiated and differentiated Caco-2 cells were infected with *C. difficile* spores previously incubated with 1, 2.5, 5, 10 and 25µg mL^−1^ of human purified E-cadherin (5085, Sigma-Aldrich, USA) in DMEM or DMEM without FBS as control for 1 h at 37 °C. Then cells were infected and were incubated for 3 h at 37 °C. Cells were washed two times with PBS to remove unbound spores, fixed with PBS–4% paraformaldehyde for 10 min at RT, and adhered spores were stained as described below.

To evaluate whether the presence of E-cadherin decreases the spore adherence in TcdA/TcdB intoxicated cells, undifferentiated and differentiated Caco-2 cells were 8 h-TcdA/TcdB intoxicated and were infected for 1 h at 37 °C with *C. difficile* spores previously incubated with 25 µg mL^−1^ of E-cadherin for 1 h at 37 °C. Cells were washed two times with PBS to remove unbound spores, fixed with PBS–4% paraformaldehyde for 10 min at RT, and adhered spores were stained as described below.

In undifferentiated, differentiated, and polarized *C. difficile* spore infected Caco-2 cells, adhered *C. difficile* spores were immunostained using the intracellular spore exclusion immunostaining assay as previously described (20). For this *C. difficile* spore–infected cells were washed three times with PBS at RT, blocked with PBS–3% BSA overnight at 4 °C, and samples were incubated with anti–*C. difficile* spore goat serum 1:50 (PAC 5573, Pacific Immunology, USA) in PBS–1% BSA for 1 h at RT. The cells were washed 3 times with PBS and incubated with 1:400 secondary antibody donkey IgG anti–goat CFL 488 (SC362255, Santa Cruz Biotechnology, USA) for 1 h at RT. Cells were washed, and DNA was stained with 8 µg mL^−1^ Hoechst 33342 (ThermoFisher, USA) for 15 min at RT. In the case of polarized cells, to stain the F–actin, samples were incubated with 1:100 Phalloidin Alexa Fluor 568 (A12380, Invitrogen, USA) for 30 min at RT. Samples were washed three times with PBS and one time with distilled water and mounted with Dako Fluorescent Mounting Medium (Dako, Denmark). Samples were visualized in the epifluorescence microscope Olympus BX53 fluorescence microscopy with UPLFLN 100× oil objective (numerical aperture 1.30). Images were captured with the camera for fluorescence imaging Qimaging R6 Retiga, and pictures were analyzed with ImageJ (NIH, USA). Adhered spores were considered as spores in phase–contrast that were marked in fluorescence. Internalized spores were considered spores in phase–contrast, but that do not have fluorescence. In the case of polarized cells, it was only possible to quantify adhered spores. The number or adhered spores were quantified, and data were normalized to the control.

### ELISA of E-cadherin bound to *C. difficile* spores

Ninety six-well plate were coated overnight at 4ºC with 100 µL/well containing 1.6 × 10^8^ spores mL^−1^ or with 100 µL of PBS containing 1 µg of BSA in PBS (as a control). overnight at 4 °C. Wells were washed 5 times with PBS–0.05% Tween-20 (PBS–Tween) to remove unbound spores and blocked with 200 µL of PBS–Tween–2% BSA (Sigma Aldrich, USA) for 1 h at 37 °C. The spores were washed and incubated with 50 µL of 0.2, 2.0, 20, and 200 nM of recombinant human purified E-cadherin (5085, Sigma-Aldrich, USA) for 1 h at 37 °C. Unbonded E-cadherin was washed 5 times with 150 µL PBS–Tween and was incubated with 50 µL of 1:50 IgG of mouse anti–E-cadherin (SC8426-Santa Cruz Biotechnology, USA) in PBS–Tween–1% BSA for 1 h at 37 °C. Wells were washed five times with PBS–Tween and then incubated with 50 µL of 1: 5,000 secondary antibody anti–mouse-HRP (610-1302, Rockland, USA) 1 h at 37 °C. Then, wells were washed five times with PBS–Tween and once with carbonate buffer (pH 9.6). Finally, wells were incubated with 50 µL of buffer substrate (17 mM citric acid, 65 mM potassium phosphate, 0.15% hydrogen peroxide, and 0.4% θ-phenylenediamine) for 20 min at RT. The reaction was stopped with 25 μL of 4.5 N H_2_SO_4_. The color intensity was quantified at 450 nm using the plate reader Infinite F50, (Tecan, Switzerland).

### Immunofluorescence of E-cadherin bound to *C. difficile* spores

The immunofluorescence was performed as previously described with modification (19). Briefly, 4 × 10^7^ *C. difficile* spores were incubated with 1, 2.5, 5.0, 10, and 25 µg mL^−1^ of recombinant human purified E-cadherin (5085, Sigma-Aldrich, USA) in PBS for 1 h at 37 °C. Spores were washed 5 times with PBS and fixed with PBS–4% paraformaldehyde in poly–L-lysine-treated 12 mm glass–coverslips (Marienfeld GmbH, Germany). The spores were washed 3 times with PBS and were blocked with PBS–3% BSA for 1 h at RT. Then, samples were incubated with 1:50 mouse IgG anti–E-cadherin (SC8426, Santa Cruz Biotechnology, USA) for 2 h at RT and washed 3 times with PBS and once with distilled H_2_O. Finally, samples were mounted with Dako Fluorescent Mounting Medium (Dako, Denmark), and images were captured in an epifluorescence microscope Olympus BX53. The fluorescence intensity was quantified using Fiji (ImageJ, NHI, USA). Data were normalized as (Fl. Int/area)_sample_ – (Fl. Int/area)_background._ The Surface Plot was generated with the plug–in 3D surface Plot of Fiji (ImageJ, NHI, USA).

### SDS-PAGE and western blot of intoxicated Caco-2 cells

Undifferentiated and differentiated Caco-2 cells were intoxicated for 3, 6 and 8 h with TcdA/TcdB in 24-well plates. The cells were washed and frozen at −80 °C. The frozen cells were lysed with 80 µL/well of cold RIPA buffer (50 mM Tris-HCl; 150 mM NaCl; 0.5% Deoxycholate; 1% NPO_4_; 1mM EGTA; 1mM EDTA; 0.5% SDS; protease inhibitor) in ice and centrifuged at 18,400×*g* for 15 min at 4 °C. The supernatant was collected and 20 µg of protein was suspended in 2× SDS-PAGE sample loading buffer, boiled and electrophoresed on 12% and 4% acrylamide SDS-PAGE gels. Proteins were transferred to nitrocellulose membrane and blocked with 5% non-fat milk in TBS for 1 h at RT. Membranes were probed for non-glucosylated Rac1 in Tris -buffered saline containing 0.1% Tween (TTBS) with 5% milk and 1:1,000 mouse anti-Rac1 (Clone 102/Rac1 610650; BD Bioscience, USA) and then incubated overnight at RT and 6 h at RT. The membranes were washed with TTBS and incubated with 1:5,000 goat anti-mouse HRP for 1 h at RT (Rockland 610-1302) in 5% milk-TTBS. Samples were washed 3 times with TTBS and HRP activity was detected with a chemiluminescence detection system (Li-core) using PicoMax sensitive chemiluminescence HRP substrate (Biorad, USA). To detect total Rac-1 in undifferentiated cells, membranes were incubated twice with mild stripping solution (1.5% glycine, 0.1% SDS, 1% Tween 20; pH 2.2) for 7 min at RT, washed twice with PBS and twice with TTBS 0.05%, and blocked with 5% non-fat milk-TBS 1 h at RT. For differentiated cells, total Rac1 was detected in the other membrane loaded simultaneously with the same concentration of protein. Membranes were incubated with 1:1,000 mouse anti-Rac1 (clone 23A8 05-389; Merck Millipore, USA) overnight at 4 °C and 3 h at RT and then washed 3 times with TTBS. Membranes were then incubated with 1:5,000 goat anti-mouse HRP (Rockland 610-1302) 1 h at RT and washed 3 times with TTBS. HRP activity was detected using Li-core as described for non-glucosylated Rac1.

### Transmission electron microscopy of E-cadherin bound to *C. difficile* spores

2.5 × 10^8^ *C. difficile* spores were incubated with 10 µg mL^−1^ of recombinant human purified E-cadherin (5085, Sigma-Aldrich, USA) with PBS for 1 h at 37 °C. Spores were washed 5 times with PBS and centrifugation at 18,400×*g* for 5 min and were blocked with PBS–1% BSA for 20 min at RT. Then samples were incubated with 1:50 IgG of mouse anti–E-cadherin (SC8426, Santa Cruz Biotechnologies, USA) in PBS–1% BSA for 1 h at RT, washed 3 times by centrifugation at 18,400×*g* for 5 min with PBS. Spores were incubated with IgG anti–mouse conjugated to 10nm gold nanoparticles (G7777, Sigma-Aldrich, USA) for 1 h at RT. Then were washed 3 times with PBS–0.1% BSA by centrifugation at 18,400×*g* for 5 min. Subsequently, spores were processes for TEM, as previously have been described (19). Briefly, spores were fixed in 0.1 M cacodylate buffer–2.5% glutaraldehyde with 1% paraformaldehyde and then fixed with 0.1 M cacodylate buffer–1% osmium tetroxide and stained with 1% tannic acid for 30 min. Samples were dehydrated with increasing acetone concentrations; 30% acetone (with 2% uranyl acetate only in this step) for 30 min, 50% acetone for 30 min, 70% acetone overnight, and 90% acetone for 30 min, and 2 times with 100% acetone. Dehydrated samples were embedded in resin: acetone in a ratio of 3:1, 1:1, and 1: 3 for 40 min each. Finally, samples were resuspended in resin for 4 h and incubated for 12 h at 65 °C. With a microtome, 90 nm sections cut and placed in a carbon–coated grid for negative staining and double staining with 2% uranyl acetate with lead citrate. Sections were observed in a transmission electron microscope Phillip Tecnai 12 bioTwin of the Pontificia Universidad Católica de Chile. The distribution of thin or thick exosporium was quantified according to previously described morphotypes (34).

## RESULTS

### Characterization of accessible E-cadherin in monolayers of Caco-2 cells

Caco-2 cells have been employed to assess *C. difficile* interaction with IECs (18-20, 32, 33, 35-37). Early work demonstrated that *C. difficile* spores had a tropism towards the cell-cell junction of EDTA-treated Caco-2 cells (18). Therefore, to understand the dynamic of adherens junction closure during Caco-2 cell differentiation, we assessed the accessibility of E-cadherin in undifferentiated and differentiated monolayers of Caco-2 cells. First, Caco-2 cell differentiation was assessed for the appearance of microvilli by TEM and sucrase-isomaltose as a differentiation marker in monolayers cultured for 2- or 8-days post-confluency (19). Next, differentiated and undifferentiated Caco-2 cells were immunostained for accessible E-cadherin or permeabilized and stained for total E-cadherin (Figure 1*A, B*). Results demonstrate that in undifferentiated Caco-2 cells, ∼84% of the cells were immunolabeled for accessible E-cadherin (Figure 1*C*), whereas, in differentiated Caco-2 cells, only ∼7% of the cells were stained for accessible E-cadherin (Figure 1*C*). As a control, cells were permeabilized and stained for total E-cadherin, and we observed that ∼100% of both undifferentiated and differentiated Caco-2 cells were immunolabeled for E-cadherin (Figure 1*D*-*F*). These results demonstrate that after 8-days of differentiation, most of the cells lack accessible E-cadherin.

**Figure 1.**
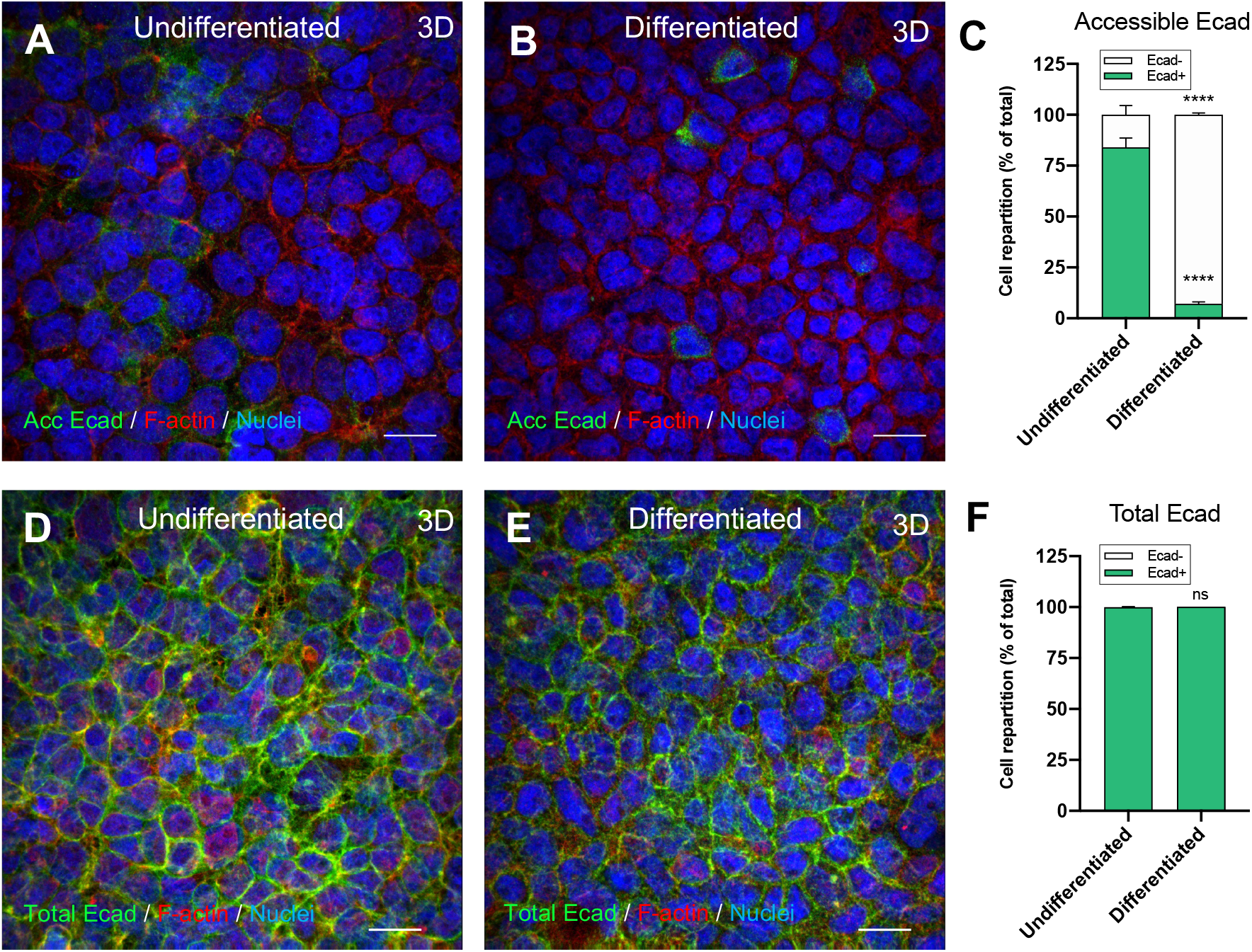
Distribution of accessible and total E-cadherin in undifferentiated and differentiated Caco-2 cells. 3D projection of a representative confocal micrograph of healthy *A*, undifferentiated, and *B*, differentiated Caco-2 cells immunostaining for accessible E-cadherin. Accessible E-cadherin (shown as acc Ecad) is in green, F-actin in red, and nuclei in blue. *C*, Repartition of cells that were positive (Ecad+) or negative (Ecad−) stained for accessible E-cadherin. 3D projection of a representative confocal micrograph of a healthy *D*, undifferentiated and *E*, differentiated Caco-2 cells immunostaining for total E-cadherin and *F*, Repartition of cells that were positive (Ecad+) or negative (Ecad−) stained for total E-cadherin. Fluorophores colors were digitally reassigned for a better representation. Micrographs are representative of 6 fields of 2 independent experiments. Scale bar, 20μm. Error bars indicate mean ± SEM. Statistical analysis was performed by two-tailed unpaired Student’s *t*-test, ns, *p* > 0.05; ****, *p* ≤ 0.0001.

### Intoxication with *C. difficile* TcdA and TcdB increases accessible E-cadherin

*C. difficile* toxins TcdA and TcdB cause tight and adherens junction dissociation (9-11). TcdA, but not TcdB, causes a displacement of E-cadherin from the cell-cell junctions to be located around the cells in Caco-2 cells (11) while in colonic epithelium, TcdB causes adherens junctions disruption (10). Since most of the *C. difficile* strains causing disease produces TcdA and TcdB (38), we asked whether TcdA and TcdB affect the accessibility of E-cadherin in differentiated Caco-2 cells. TcdA/TcdB-intoxicated differentiated Caco-2 cells for 3, 6, and 8 h were immunostained for accessible E-cadherin (non-permeabilized) using an antibody that detects an extracellular domain of E-cadherin, followed by fluorescent-labeled secondary antibody. For total E-cadherin staining, cells were permeabilized and immunostained with the same primary antibody and a fluorescent-labeled secondary antibody with another Alexa Fluor dye (See Methods). Then, we acquired confocal micrographs, and the fluorescence intensity (Fl. Int.) was quantified for every slice of the *z*-stack for accessible and total E-cadherin. The sum of the Fl. Int. of accessible E-cadherin was significantly higher (175% relative to unintoxicated monolayers) after 8h-intoxication (Figure 2*A, B*, Supplementary Figure 1). To evaluate if this increase of Fl. Int. occurs in the apical or basal half of the monolayer, we split the number of slices of the z-stack into 2-halves (apical and basal) with the same number of slices. We summed the Fl. Int. for accessible E-cadherin in the apical (Figure 2*C*) or basal (Figure 2*D*) halves of the monolayer. We observed a significant increase in accessible E-cadherin Fl.Int. in both the apical and basal halves, 177% and 176% respectively, after 8h-intoxication (Figure 2*C, D*). Using the same strategy, we quantified the total E-cadherin Fl. Int. in the complete *z*-stack, and we observed a significant decrease of 19% after 6h- and an increase of 12.2% after 8h-intoxication compared to unintoxicated cells (Figure 2*E*). Then, we observed that these variations were in the apical and not in the basal half of the monolayer. We observed a decrease of 35% after 6h- and an increase of 44% after 8h-intoxication compared to unintoxicated cells (Figure 2*F, G*). To confirm the degree of TcdA/TcdB intoxication, we evaluated the glucosyltransferase activity by measuring the levels of glucosylated Rac1 in TcdA/TcdB intoxicated differentiated and undifferentiated monolayers using strategies previously described (39). After 3 h of intoxication no modification of Rac1 was observed. However, after 6 and 8 h of intoxication, the unglucosylated Rac1 signal disappeared, indicating a complete glucosylation of Rac1 (Figure 2*H*). These results indicate that the cells were fully intoxicated by 6 hours and accessible E-cadherin is increased in differentiated Caco-2 cells after 8h-intoxication.

**Figure 2.**
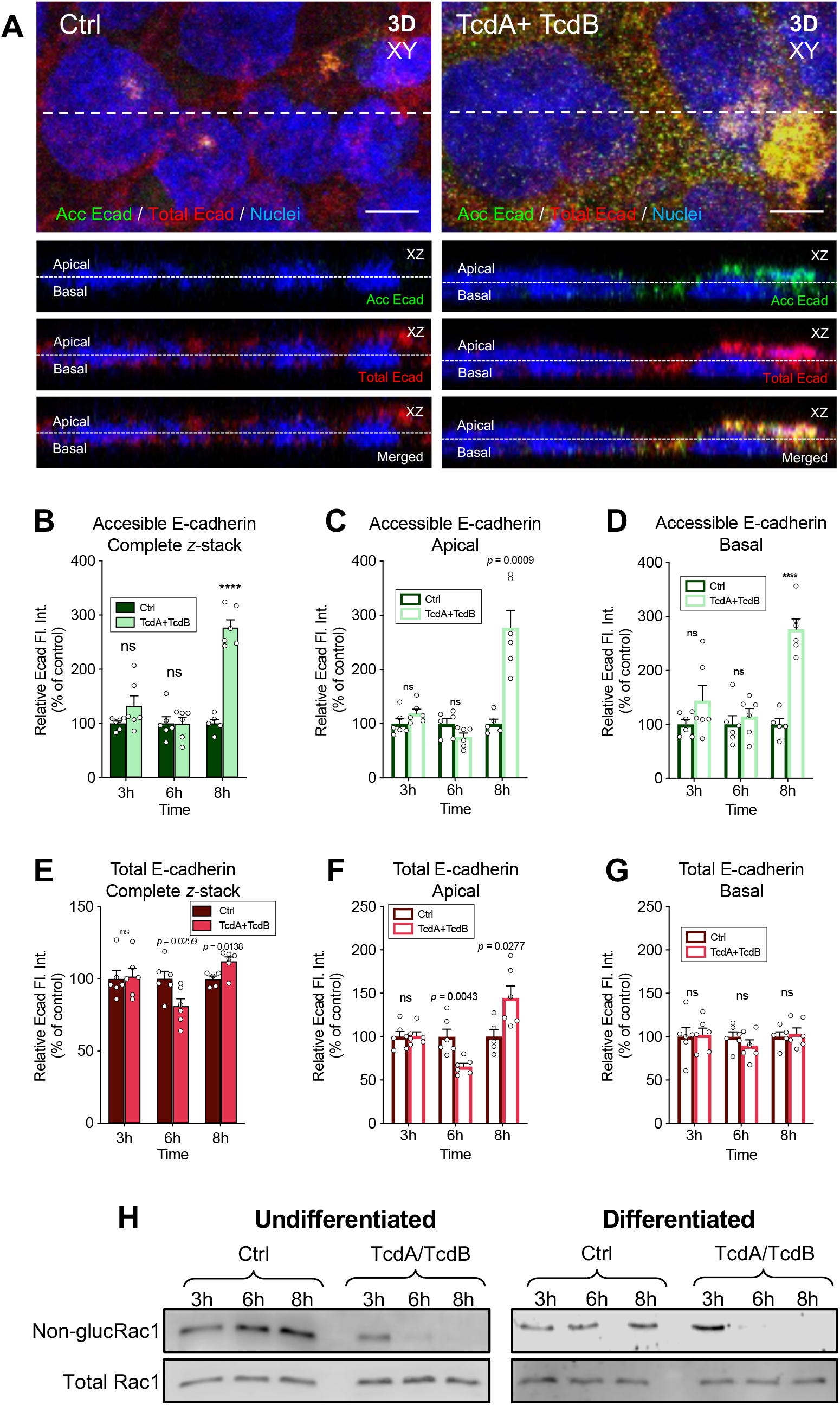
The accessibility of E-cadherin increases in TcdA and TcdB intoxicated IECs. Differentiated TcdA/TcdB intoxicated Caco-2 cells for 3, 6, or 8 h in DMEM without serum. As a control, cells were incubated with DMEM without serum. Non-permeabilized cells were stained for accessible E-cadherin (shown as acc Ecad; green), permeabilized, and total E-cadherin was stained (shown as total Ecad; red) and nuclei (blue). *A*, Representative confocal microscopy images 3D (XY) projection of control cells (left) and TcdA/TcdB intoxicated cells for 8h (right); below the orthogonal view (XZ; accessible E-cadherin, total E-cadherin and merged) of the dotted line of the XY. Representative confocal images for 3h and 8h not shown. Relative Fl. Int. measured as the sum of each z-step of *B*, accessible E-cadherin, and its abundance in the *C*, apical side, or *D* in the basal side of the cell. Relative Fl.Int. of *E*, total E-cadherin, its abundance in the *F*, apical side, and the *G*, basal side of the cell. Controls were set 100%. Error bars indicate the mean ± SEM from at least 9 fields (*n* = 3). *H*, immunoblotting anti non-glucosylated Rac1 and total Rac1 of cell lysates of undifferentiated and differentiated Caco-2 cells intoxicated with TcdA/TcdB for 3, 6 or 8h. For undifferentiated in non-glucosylated Rac1 was evaluated, then stripped and membranes were tested for total Rac1. Meanwhile for differentiated cells, the same sample was solved in 2 independent membranes simultaneously and were independently tested for non-glucosylated and total Rac1. Western blot are representative of 3 independent experiment. Statistical analysis was performed by two-tailed unpaired Student’s *t*-test, ns, *p* > 0.05; ****, *p* < 0.0001. Bars, 5μm.

### *C. difficile* toxins TcdA and TcdB increase the adherence to IECs

Several studies have shown that vegetative *C. difficile*-adherence to IECs is enhanced following TcdA or CDT intoxication (11, 40, 41). To expand the understanding of how intoxication of IECs impacts spore-adherence, differentiated and undifferentiated Caco-2 cells were TcdA/TcdB intoxicated for 8 h, or incubated 8 h with heat inactivated TcdA/TcdB as a control. Then cells were washed and infected for 1 h with *C. difficile* spores in the absence of serum. Cells were fixed and immunostained without permeabilization. In TcdA/TcdB intoxicated Caco-2 cells, we observed more adhered spores compared to unintoxicated cells (Figure 3*A*). After quantitative analysis, we observed in undifferentiated Caco-2 cells that spore adherence in 8h-TcdA/TcdB-intoxicated cells was ∼105% higher than in unintoxicated monolayers, and no differences in spore adherence were observed in cells 8 h-incubated with heat-inactivated TcdA/TcdB (Figure 3*B*). Similarly, in differentiated cells we observed an increase in ∼321% in 8 h-TcdA/TcdB-intoxicated cells over that in unintoxicated monolayers, and no differences were observed in cells 8 h-incubated with heat-inactivated TcdA/TcdB (Figure 3*C*). Collectively these data indicate that TcdA/TcdB intoxication of undifferentiated and differentiated Caco-2 cells increases *C. difficile* spore adherence.

**Figure 3.**
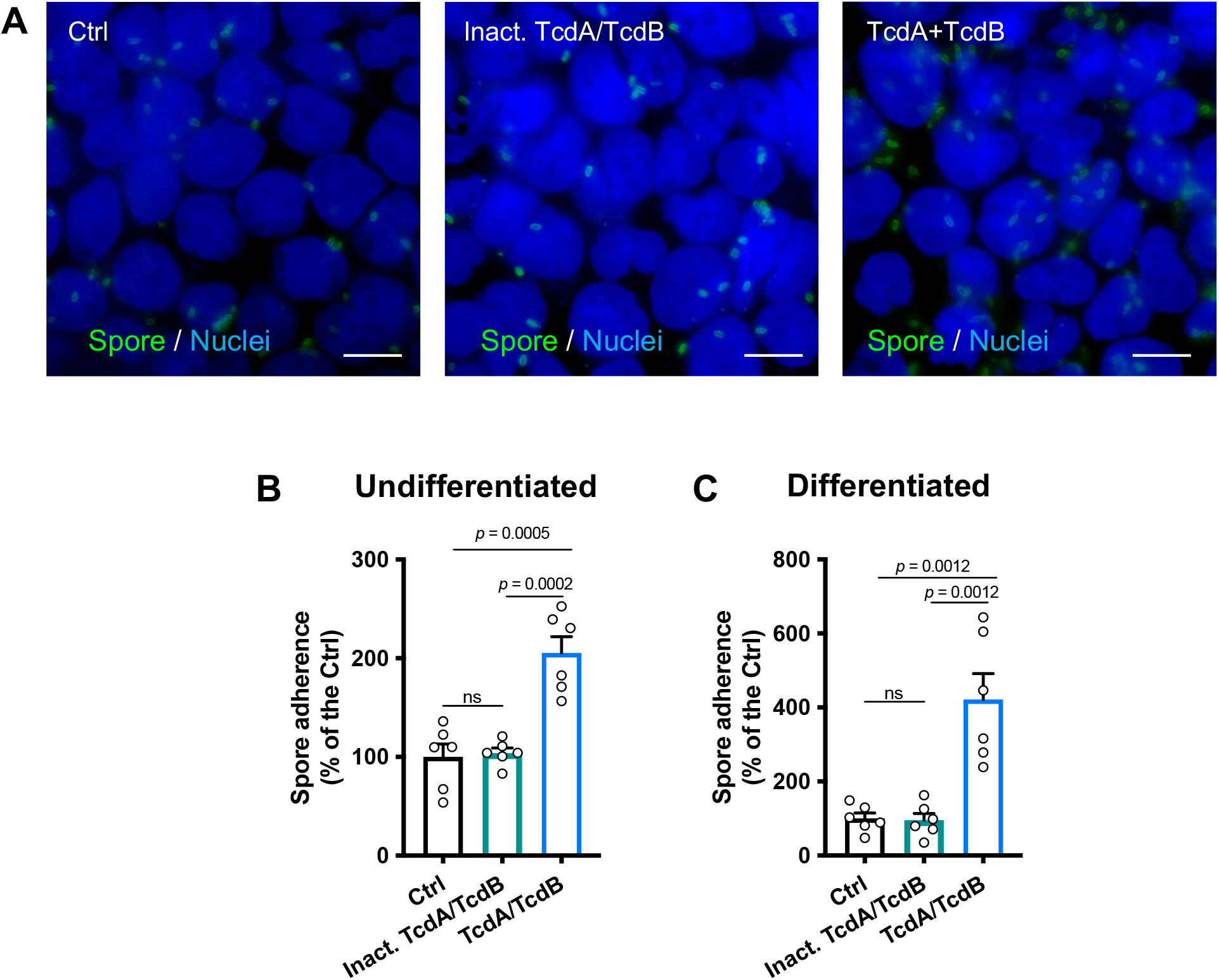
TcdA and TcdB increase *C. difficile* spore interaction with IECs. Differentiated TcdA/TcdB intoxicated Caco-2 cells for 8h or incubated with DMEM without serum as control and then, were washed with PBS and infected for 3h at an MOI 10 with *C. difficile* spores. Unbound spores were rinsed off. Non-permeabilized cells were stained for adhered spores. *A*, Representative epifluorescence microscopy images of TcdA and TcdB intoxicated cells (or DMEM as control) for 8h then washed and infected with *C. difficile* spore. *B, C. difficile* spore adherence relative in *B*, undifferentiated and *C*, differentiated cells. In bars, each dot represents one independent well from 2 independent experiment, Per well, a total of 5 fields were analyzed. Error bars indicate the mean ± SEM.. Statistical analysis was performed by unpaired *t*-test, ns, *p* > 0.05. Bars, 10 μm.

### Intoxication of IECs increases *C. difficile* spore association with E-cadherin

Since we observed that TcdA/TcdB-intoxicated differentiated Caco-2 cells lead to an increase of accessible E-cadherin and adherence of *C. difficile* spores, we tested whether this could be attributed to *C. difficile* spores associating with accessible E-cadherin. For this, unintoxicated or TcdA/TcdB-intoxicated differentiated Caco-2 cells were infected with *C. difficile* spores for 1 h. Then, accessible E-cadherin and spores were immunostained and visualized by confocal microscopy (Figure 4*A, D*). Plot profile analysis of unintoxicated Caco-2 cells (Figure 4*A*) reveals that ∼21% of adhered *C. difficile* spores were associated with accessible E-cadherin (Figure 4*B, C*, and *G*). Notably, upon TcdA/TcdB-intoxication (Figure 4*D*), ∼72% of adhered spores were associated with accessible E-cadherin (Figure 4*E, F, G*). These results indicate that adhered *C. difficile* spores can associate with accessible E-cadherin in IECs, and this association is enhanced upon TcdA/TcdB-intoxication.

**Figure 4.**
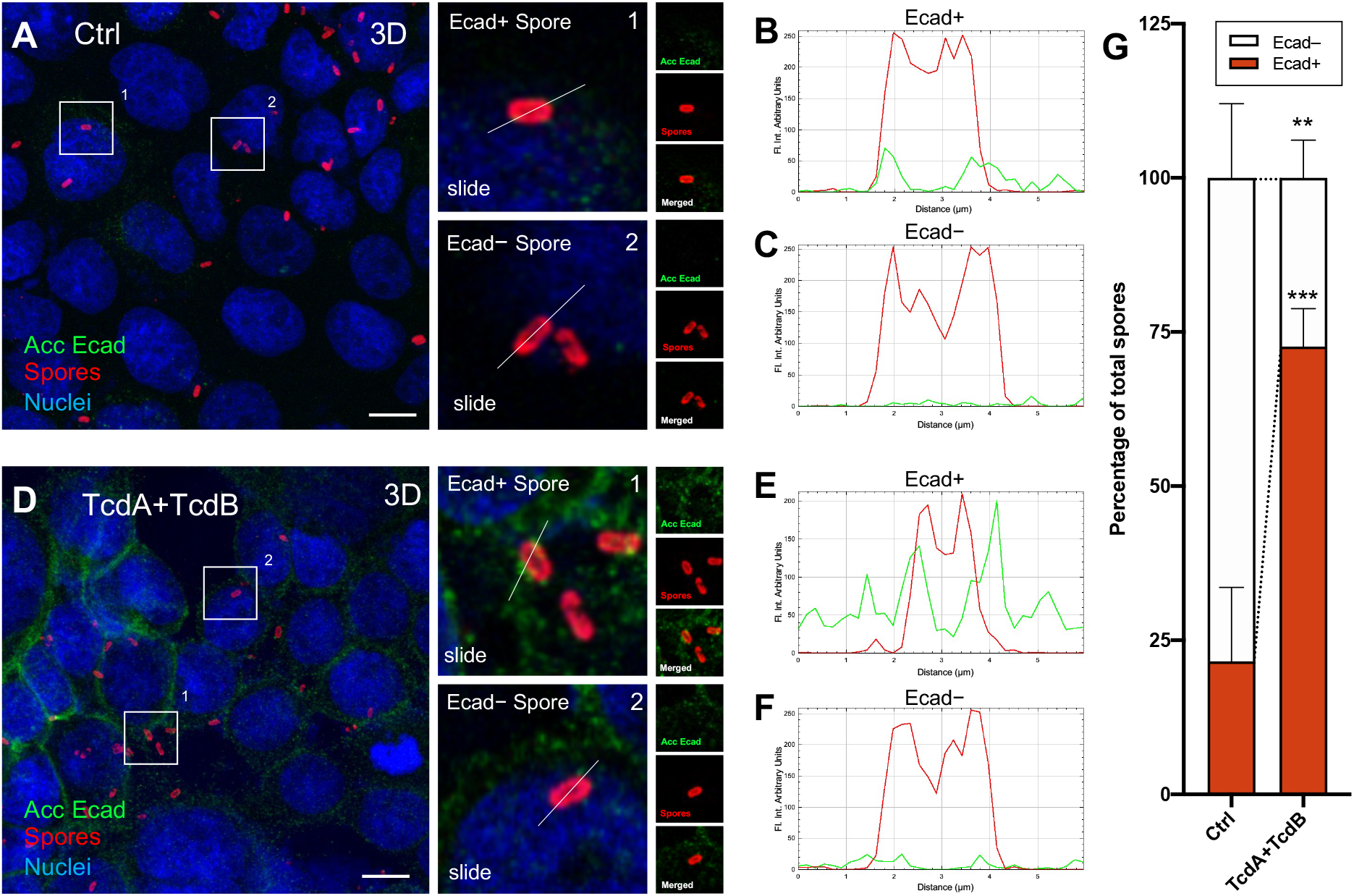
E-cadherin of IECs interacts with *C. difficile* spores increasing in TcdA and TcdB intoxicated cells. Differentiated Caco-2 cells were intoxicated with TcdA and TcdB for 8 h at 37 °C in DMEM without serum. As a control, cells were incubated with DMEM without serum. Then cells were infected with *C. difficile* spores for 1 h at 37 °C. Non-permeabilized cells were stained for accessible E-cadherin (shown as acc Ecad; green), spores (red), and nuclei (blue). *A*, Representative 3D projection confocal micrograph of healthy cells infected with *C. difficile* spores (Ctrl). On the right, magnifications of representative *C. difficile* spores associated or not with E-cadherin. *B, C*, Plot profile Fl.Int. of *C. difficile* spores (red line) and accessible E-cadherin (shown as Ecad, green line) performed in the white line of *A. D*, Representative 3D projection confocal micrograph of TcdA and TcdB intoxicated cells infected with *C. difficile* spores. On the right magnifications of representative *C. difficile* spores associated or not with E-cadherin. *E, F*, Plot profile Fl.Int. of *C. difficile* spores (red) and accessible E-cadherin (shown as Ecad, in green) performed in the white line of *D*. Repartition of spores that were positive (Ecad+) or negative (Ecad−) associated with fluorescence signal for accessible E-cadherin was shown as the average associated/non-associated spores with E-cadherin for each field, a total of ∼500 spores were analyzed. Fluorophores were digitally reassigned for a better representation. Micrographs are representative of 6 independent fields of 3 different experiments. Error bars indicate mean ± SEM. Statistical analysis was performed by two-tailed unpaired Student’s *t*-test, ns, *p* > 0.05; **, *p* ≤ 0.01; ***, *p* ≤ 0.001. Bars, 10μm.

### Antibody blocking of accessible E-cadherin reduces adherence and internalization of *C. difficile* spores to IECs

Next, we assessed whether E-cadherin is involved in *C. difficile* spore adherence and internalization. We have shown that *C. difficile* spore internalization occurs in the presence of serum (20). Therefore, undifferentiated, differentiated, and polarized Caco-2 cells were incubated with anti-E-cadherin antibody for 2h and subsequently infected for 3h with FBS-incubated *C. difficile* spores. To identify intracellular spores, we used the intracellular spore exclusion immunostaining assay (20), where only extracellular spores are immunostained in non-permeabilized cells, and total spores are visualized by phase-contrast microscopy. Using this strategy in undifferentiated Caco-2 cells, we observed that *C. difficile* spore adherence to IECs was significantly decreased upon increasing concentrations of anti-E-cadherin antibody; spore adherence decreased by 38, 53, and 73% compared to the control when cells were incubated with 1, 10, and 100 µg mL^−1^ of anti-E-cadherin antibody, respectively (Figure 5*A*). Upon quantifying internalization in undifferentiated Caco-2 cells, we observed a significant decrease of 38, 43, and 69% compared to the control in cells incubated with 1, 10, 100 µg mL^−1^ of anti-E-cadherin, respectively (Figure 5*B*). Similarly, in differentiated Caco-2 cells, a significant decrease in spore adherence in 18, 32, 60, and 70% compared to the control was observed in cells incubated with 0.1, 1, 10, and 100 µg mL^−1^ of anti-E-cadherin, respectively (Figure 5*C*). For internalization, we observed a significant decrease in 15, 35, 42, and 54% compared to the control in cells incubated with 0.1, 1, 10, and 100 µg mL^−1^ of anti-E-cadherin, respectively (Figure 5*D*). Finally, in polarized Caco-2 cells, we observed a decrease in 32, 56, 63, and 79% in spore adherence compared to the control in presence with 0.1, 1, 10, and 100 µg mL^−1^ of anti-E-cadherin, respectively (Figure 5*E*). These data indicate that the antibody-mediated blockage of accessible E-cadherin reduces *C. difficile* spore adherence and internalization to IECs.

**Figure 5.**
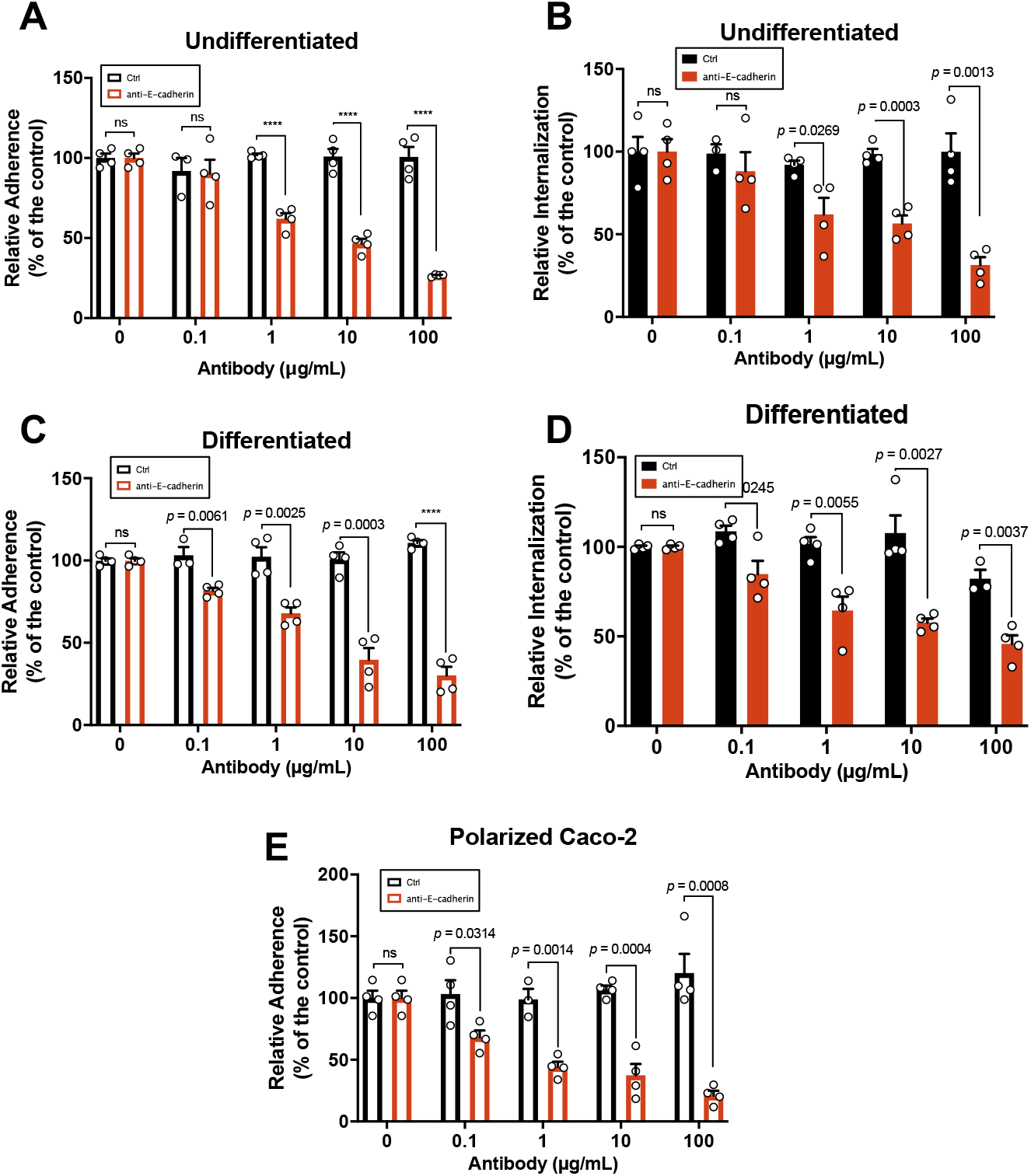
E-cadherin blocking antibodies decrease the spore interaction with IECs. Undifferentiated, differentiated, and polarized Caco-2 cells were infected with *C. difficile* spores preincubated with FBS in the presence of anti-E-cadherin or non-immune IgG as control for 2 h at 37 °C. *A*, adherence, and *B*, internalization of *C. difficile* spores in undifferentiated Caco-2 cells or *C*, adherence and *D*, internalization in differentiated Caco-2. *E, C. difficile* spore adherence in polarized Caco-2 cells. Data showed in each panel are normalized to 0 µg mL^−1^ of antibody. The data are the mean of 6 independent wells collected in 2 independent experiments. In bars, each dot represents one independent well collected from 2 independent experiments. For each bar, a total of ∼1,300 spores were analyzed. Error bars indicate mean ± SEM. Statistical analysis was performed by two-tailed unpaired Student’s *t*-test, ns, *p* > 0.05; ****, *p* < 0.0001.

### Purified recombinant human E-cadherin competitively inhibits spore adherence in undifferentiated, differentiated and TcdA/TcdB intoxicated IECs cells

Subsequently, to provide further evidence that E-cadherin acts as a spore adherence receptors, we evaluated whether infected healthy undifferentiated Caco-2 monolayers with *C. difficile* spores preincubated with 1, 2.5, 5, 10, and 25 µg mL^−1^ of E-cadherin, and we observed a decrease in spore adherence of 12%, 26%, 38%, 47% and 64% respectively compared to spores not incubated with E-cadherin (Fig. 6*E*). In differentiated Caco-2 cells, we observed a decrease in spore adherence of 27%, 30%, 30%, 27% and 53% for spores incubated with 1, 2.5, 5, 10, and 25 µg mL^−1^ of E-cadherin, respectively (Figure. 6*F*). Finally, we also assessed whether the increased spore adherence that we previously observe in TcdA/TcdB intoxicated Caco-2 cells was E-cadherin specific. As expected, significant *C. difficile* spore adherence to TcdA/TcdB intoxicated Caco-2 cells was observed in the absence of E-cadherin; however, upon of TcdA/TcdB intoxicated differentiated and undifferentiated Caco-2 cells with spores previously incubated with 25µg mL^−1^ of E-cadherin for 1 h at 37°C, and we observed a significant decrease of 41% in undifferentiated (Figure 6*G*) and 55% in differentiated Caco-2 cells (Figure 6*H*) compared to TcdA/TcdB intoxicated cells. Overall, these results show that E-cadherin interacts with *C. difficile* spores in a dose-dependent manner and that E-cadherin-ligand binding sites are non-homogeneously distributed on the *C. difficile* spore surface. There results indicate that competition experiments with purified E-cadherin reduces the *C. difficile* spore adherence in healthy and intoxicated cells.

**Figure 6.**
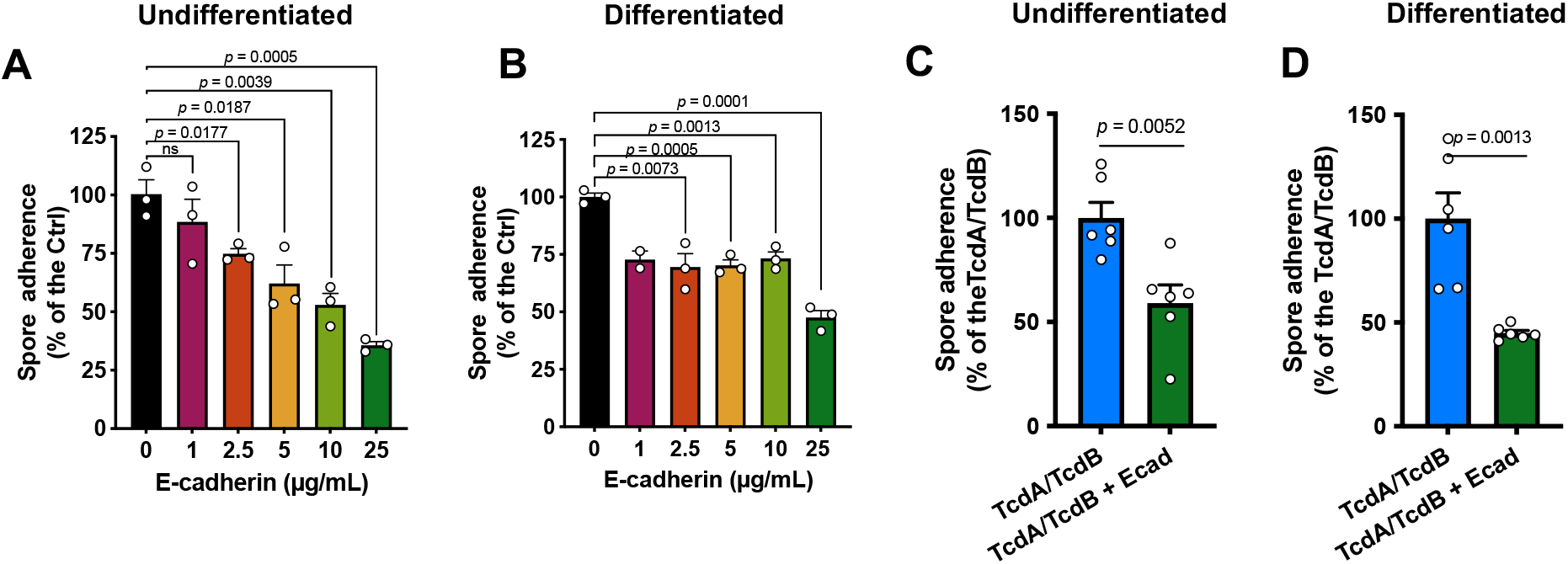
E-cadherin competitively reduces spore adherence to healthy and intoxicated IECs. *A* Undifferentiated and differentiated Caco-2 cells where infected for 3 h at a MOI of 10 with *C. difficile* spores pretreated with 1, 2.5, 5, 10 and 25µg/mL of human purified E-cadherin, and adhered cells where identified and quantified as described in the Method section. *B*. TcdA/TcdB intoxicated undifferentiated and differentiated Caco-2 cells where infected for 3 h at a MOI of 10 with *C. difficile* spores pretreated with 25µg/mL of human purified E-cadherin, and adhered cells where identified and quantified as described in the Method section. The data are the mean of 6 independent wells collected in 2 independent experiments. Adherence of *C. difficile* spores preincubated with E-cadherin (or DMEM as control) in 8 h-TcdA/TcdB intoxicated cells. The data are the mean of 6 independent wells collected in 2 independent experiments.

### E-cadherin binds in a concentration-dependent manner to *C. difficile* spores

To provide evidence that *C. difficile* spores interact with E-cadherin, we used ELISA with *C. difficile* spore-coated wells incubated with increasing concentrations of E-cadherin, or BSA as a control. E-cadherin binding activity was evident in *C. difficile* spore-coated wells (Figure. 7*A*). The specificity of this interaction was demonstrated by the dose-dependent binding. Additionally, we evaluated the distribution of binding of E-cadherin to the spore surface by immunofluorescence. Upon micrograph analysis, we observed that anti-E-cadherin immunostained spores exhibited a non-homogeneous Fl. Int surrounding the spore (Figure. 7*B*). In spores, the Fl. Int. for E-cadherin was 156, 189, 258, 304, and 369% relative to the control upon incubation with 1, 2.5, 5, 10, and 25 µg mL^−1^ of E-cadherin (Figure. 7*C*). Next, we established a Fl. Int cut-off limit for those *C. difficile* spores that were visually associated with 2.5 µg mL^−1^ E-cadherin. Based on this threshold, 59, 65, and 66% of the spores were positive for E-cadherin in the presence of 5, 10, and 25 µg mL^−1^ E-cadherin, respectively. Finally, we evaluated E-cadherin Fl. Int distribution for positive immunodetected spores, and we observed that they were mainly distributed between 1.4-2.1 normalized Fl. Int. arbitrary units (Figure. 7*D*). Taken together, these results indicate that E-cadherin binds in a concentration dependent manner to *C. difficile* spores, in a heterogeneous and skewed manner to *C. difficile* spores.

**Figure 7.**
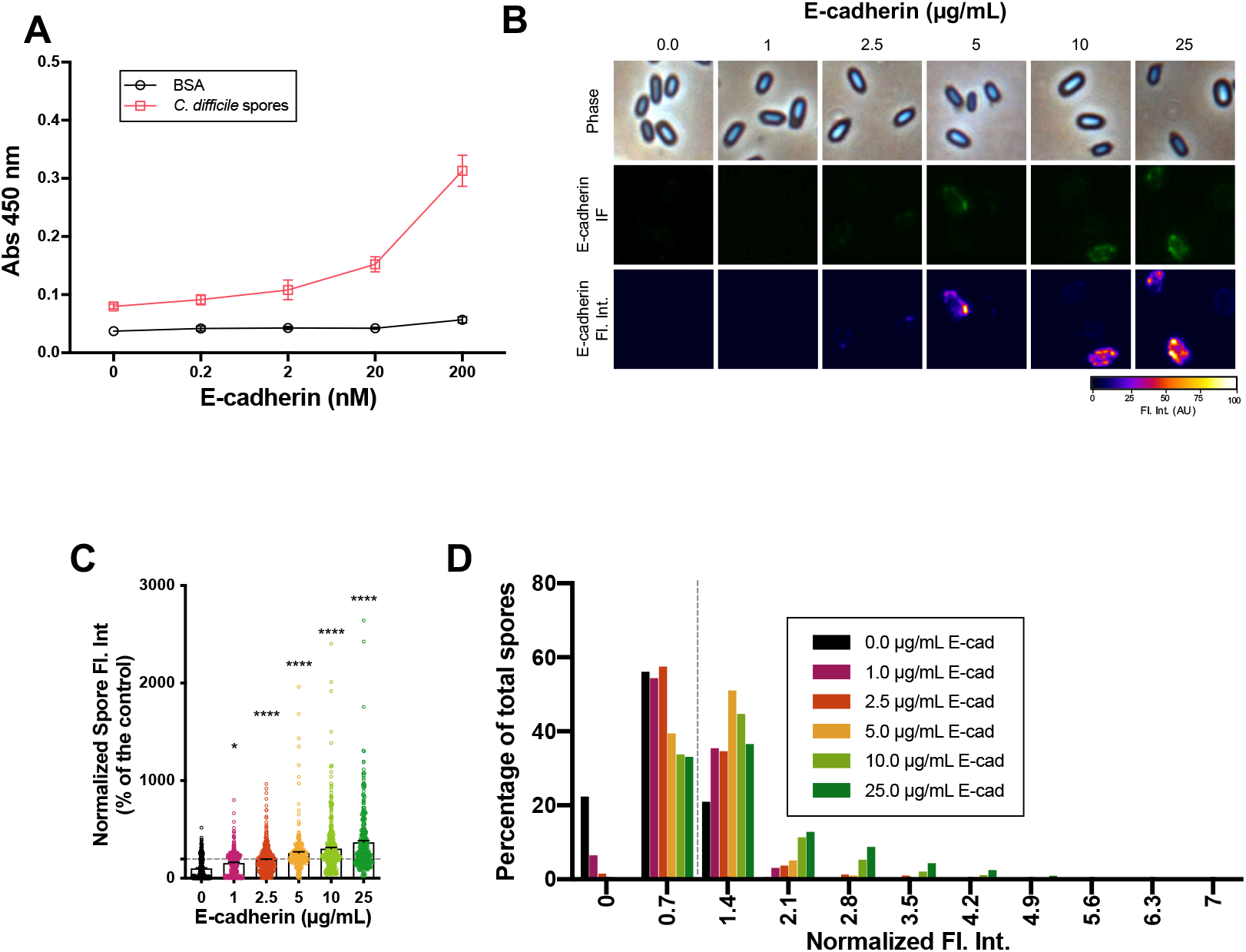
E-cadherin interacts with *C. difficile* spore. *A*, Solid-phase binding assay of *C. difficile* R20291 spores or BSA as control with increasing concentrations (i.e., 0.2, 2.0, 20 and 200 nM) of human purified E-cadherin. The data represented are the mean of 9 independent wells collected in 3 independent experiments. *B*, Representative phase-contrast micrograph, immunofluorescence against E-cadherin, and immunofluorescence plot profile of 4 × 10^7^ *C. difficile* spores that were incubated with increasing concentrations (i.e., 1, 2.5, 5, 10, 25 µg mL^−1^) of human purified E-cadherin for 1 h at 37 °C. *C*, Quantitative analysis of Fl.Int. *D*, Distribution of E-cadherin fluorescence intensity of *C. difficile* incubated with increasing concentrations (i.e., 1, 2.5, 5, 10, 25 µg mL^−1^) of human purified E-cadherin. A total of ∼450 spores were analyzed for each condition. Error bars indicate mean ± SEM. Statistical analysis was performed by two-tailed unpaired Student’s *t*-test, ns, *p* > 0.05; *, *p* < 0.05; ****, *p* < 0.0001.

### E-cadherin interacts with hair-like projections of *C. difficile* spores

The outermost structure of *C. difficile* spores are the hair-like projections, formed in part by BclA3, that are common in epidemically relevant strains (16, 20, 42). Isogenic sporulating cultures of *C. difficile* lead to the formation of spores with two exosporium morphotypes that exhibit hair-like projections, spores with a thick exosporium layer display distinctive electron-dense bumps, and spores with a thin exosporium lack these bumps (34, 42-44), suggesting that this exosporium morphotype diversity might be contributing to E-cadherin spore-binding variability. Consequently, to gain ultrastructural insight of the spore-E-cadherin interaction, *C. difficile* spores incubated with E-cadherin were imaged by TEM and immunogold-labeling of E-cadherin. We observed E-cadherin-gold particles in close contact with the exosporium of *C. difficile* spores, particularly with the hair-like projections (Figure 8*A*-*E*). No gold particles were observed in spores without the E-cadherin incubation. (Figure 8*A*). Next, within the E-cadherin positive spores, we observed gold particles in 52/236 (22.1%) of the spores (Figure 8*F*). Of these, 19/52 (36.5%) and 33/52 (63.5%) of the E-cadherin-bound spores corresponded to spores with thin and thick exosporiums, respectively (Figure 8*G*). These results are representative of a 90 nm-slice of the sample, and therefore the number of gold particles corresponds to those only present in the slice and not of a complete spore. These results suggest that E-cadherin binds preferentially to thick-exosporium layer spores, which seem to have enriched in hair-like projections. and specifically to the hair-like projections.

**Figure 8.**
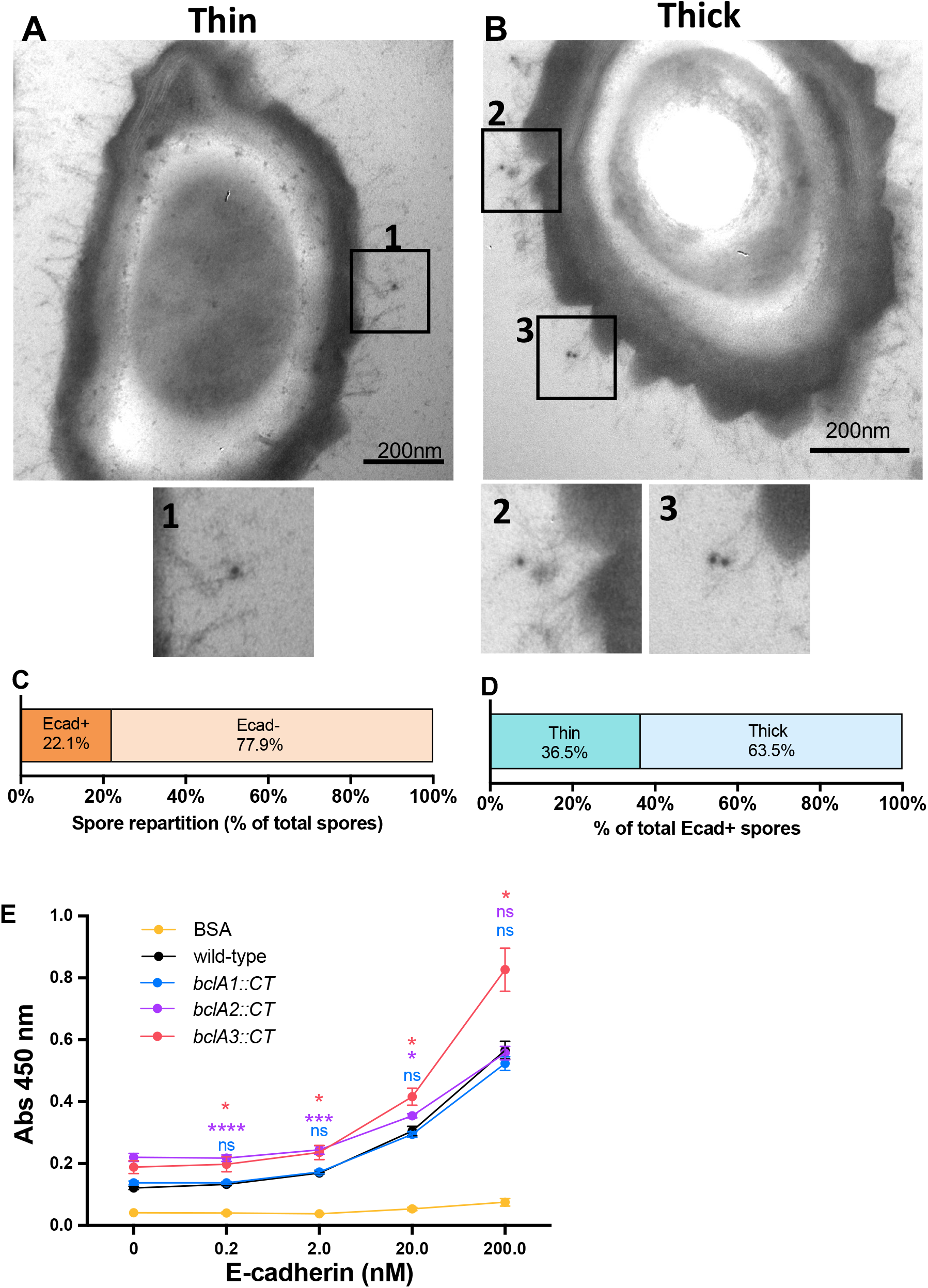
Immunoelectron microscopy of E-cadherin-gold complex binds to *C. difficile* spores. *A,B* TEM of *C. difficile* spores incubated with 10 µg mL^−1^ E-cadherin or without E-cadherin (negative control). Then incubated with anti-E-cadherin and anti-mouse-gold 12 nm complex. White arrows denote the mouse anti-E-cadherin and anti-mouse-gold 12 nm complex. Spores where stratified in thick and thin exosporium spores. Lower panels are magnification of the black squares highlighting immunogold spots, Quantification of the *C. difficile* spore associated with E-cadherin (“−” and “+” denote negative or positive association, respectively). *C,D* Classification of E-cadherin positively associated *C. difficile* spores according to the morphology of the spore, thin or thick exosporium, represented as the percentage of the total E-cadherin positive spores. Figures show representative spores with a thin or thick exosporium layer, and are representative of 2 independent experiments. A total of 100 spores were quantified. *E* solid-phase binding assay of human E-cadherin to *C. difficile* spores of 630 wild-type and isogenic mutant strains of *bclA1, bclA2* and *bclA3* or BSA as a control with increasing concentrations (i.e., 0.2, 2.0, 20 and 200 nM) of human purified E-cadherin. The data represented are the mean of 9 independent wells collected in 3 independent experiments. Statistical analysis was performed by ONE-WAY ANOVA followed by Bonferroni for multicomparisons, ns, *p* > 0.05; *, *p* < 0.05, ****, *p* < 0.0001. Colors correspond to the mutant strain being compared to wild-type.

Most *C. difficile* genomes encode three *bclA* orthologues encodes three collagen-like proteins of the BclA family of proteins (45). However, *C. difficile* strain R20291 only encodes two *bclA* orthologues (i.e., *bclA2* and *bclA3*) (45). Therefore, we reasoned to use previously characterized *bclA1, bclA2* and *bclA3* mutants in strain 630 (30) to address by ELISA whether these collagen-like proteins could be implicated in E-cadherin binding to *C. difficile* spores. As with R20291 spores, E-cadherin binding to 630 spore coated wells was evident (Figure 8E). Notably, upon coating wells with spores of *bclA1, bclA2* or *bclA3* spores, a dose dependent binding of E-cadherin to spore-coated wells that as similar to that of wild-type spores, with the exception of *bclA3* spores which exhibited higher binding of E-cadherin (Figure 8E). Overall, these results suggest the presence of an unidentified E-cadherin-specific spore surface ligand.

## DISCUSSION

A major clinical challenge that remains unresolved is how to address the high rates of CDI recurrence (46-48). *C. difficile* spores are regarded as the key virulence factors required for recurrence and transmission of the disease (16, 17). Although significant work has advanced our understanding of the biology of *C. difficile* spores (16, 42), the mechanisms through which *C. difficile* spores interact with the host within the environment of the colon remain poorly described. The integrity of the intestinal epithelial barrier is key in preventing pathogen translocation (22). However, in the healthy intestinal mucosa, the apical- and basolateral-cell sides are not always separated. The adherens junctions, where E-cadherin is located, can be accessible in certain situations, i.e., along lateral membranes of goblet cells (20, 49, 50), extruding apoptotic sites, produced transiently when senescent cells are expelled and detached from the epithelium (49, 51, 52) and in epithelial folds of the intestinal villus produced by tension and constriction forces (49). In this context, we have demonstrated that E-cadherin is a novel spore adherence-receptor that promotes binding of *C. difficile* spores at adherens junctions, and this binding is enhanced in intoxicated IECs.

Work by others have demonstrated that E-cadherin is hijacked by several pathogens, including *Listeria monocytogenes, Fusobacterium nucleatum*, and *Streptococcus pneumoniae* (23-26, 49). This report expands the list of pathogens that hijack E-cadherin for adherence and entry to include *C. difficile* spores. Our prior work had demonstrated that *C. difficile* spores adhere to IECs *in vitro* (18) and gain entry into host cells via fibronectin and vitronectin in an integrin-dependent pathway (20). The ∼70-80% reduction in adherence in undifferentiated, differentiated and polarized monolayers of Caco-2 cells upon blockade with anti-E-cadherin antibodies or competition with purified E-cadherin, suggests that E-cadherin is a major cellular adherence receptor for *C. difficile* spores. Fibronectin, vitronectin, and integrin receptors are typically located in the basolateral membrane (41, 53-56), where E-cadherin is localized. An outstanding question is whether E-cadherin acts as an anchoring protein allowing *C. difficile* spores to bind fibronectin/vitronectin and their cognate integrins to trigger internalization, or whether spore-binding to E-cadherin alone is capable of triggering a signal for internalization of *C. difficile* spores. This question will be addressed in future studies.

Another conclusion of this work is that *C. difficile* spores bind to E-cadherin in a dose-dependent manner. *L. monocytogenes* interacts with human and not with mouse E-cadherin through InlA (26). While the exosporium of *C. difficile* spores has no orthologs of InlA (57), we show E-cadherin binding to the hair-like projections of the exosporium. *C. difficile* spores form two exosporium morphotypes during sporulation (i.e., a thick and a thin exosporium layer) (42); among the main differences is the presence of electron-dense bumps underlying the hair-like projections. Whether there are differences in the hair-like projections and in their numbers between both morphotypes, is unclear, and matter of study in our laboratory. The relevance of the exosporium thickness in E-cadherin binding. Notably, spores of *bclA1* and *bclA2* orthologue in strain 630 were able to bind to E-cadherin to a similar extent as wild-type spores, suggesting either redundance in function. It was surprising to observe that *bclA3* mutant spores bind higher levels of E-cadherin, suggesting that absence of BclA3 might uncover an alternative spore surface ligand(s) implicated in binding of E-cadherin to the spore surface. Current ongoing work in our laboratory aims to identify the spore surface ligand(s) that bind to E-cadherin and its implication in disease.

A final and major conclusion of this work is that TcdA/TcdB intoxication increases spore adherence to IECs in a E-cadherin-specific manner. This is supported by increased association between basolateral E-cadherin and *C. difficile* spores, and by the observation that soluble E-cadherin can reduce the adherence of *C. difficile* spores to TcdA/TcdB intoxicated IECs. *C. difficile* toxins TcdA and TcdB cause a disruption of the cell-cell junctions (10, 12, 13), causing a loss of cellular polarity (9-11). This disruption is caused by a dissociation of tight and adherens junction proteins. In this context, our work contributes to understanding how *C. difficile* spores interact with TcdA/TcdB-intoxicated IECs. Our results demonstrate that differentiated TcdA/TcdB-intoxicated monolayers show an increased accessibility of E-cadherin that correlates with an increase in spore adherence to TcdA/TcdB-intoxicated cells. These observations suggest that during CDI, TcdA and TcdB increase the E-cadherin accessibility, contributing to the spore adherence and entry into the IECs *in vivo*. Indeed, since intracellular *C. difficile* spores contribute to disease recurrence (20), TcdA/TcdB -intoxication may significantly contribute to *C. difficile* spore adherence and internalization during CDI, contributing overall to disease recurrence. These hypotheses are in agreement with clinical observations in CDI patients where high titers of endogenous antibodies against the *C. difficile* toxins correlate with reduced R-CDI rates (58), and with the reduced R-CDI observed in the clinical trials of bezlotoxumab (15). The development of multifactorial treatment with intervention in TcdA/TcdB, spore adherence, and entry to IECs might importantly reduce the R-CDI rates.

## Supplementary figures review only

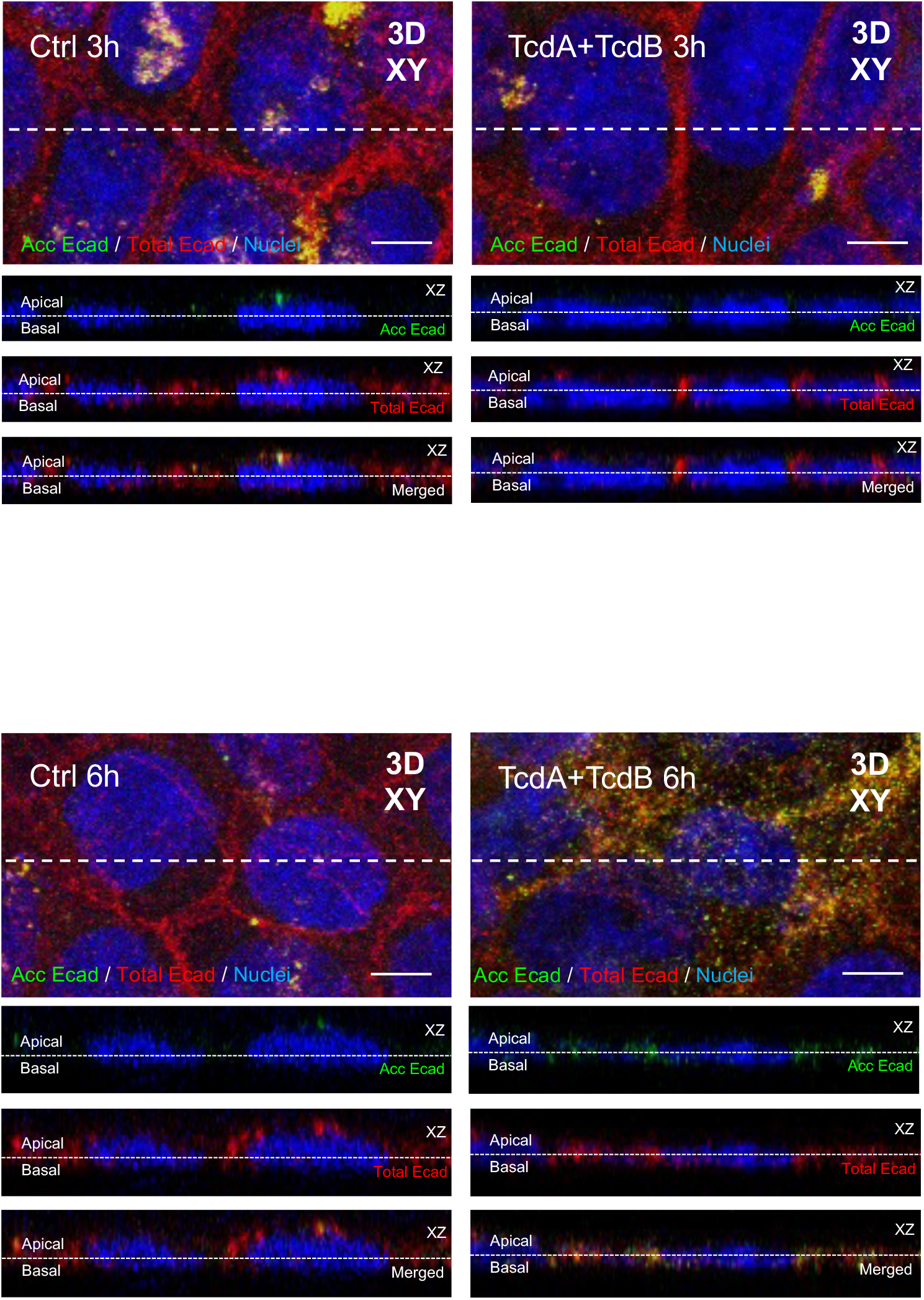
Supplementary confocal micrographs of Fig. 1. These are supplementary figures from the experiment detailed in Fig. 4. Differentiated Caco-2 cells intoxicated with 600 pM of TcdA and TcdB for 3, 6, or 8 h in DMEM without serum. As a control, cells were treated with DMEM without serum. Non-permeabilized cells were stained for accessible E-cadherin (shown as acc Ecad) in green, then permeabilized and stained total E-cadherin (total Ecad) shown in red nuclei shown in blue. Representative confocal microscopy images 3D (XY) projection of intoxicated cells for 3 h (left) or 6 h (right) with its control; below a magnified slide (XY), and the orthogonal view (XZ). Representative images from at 9 fields (*n* = 3). Bars, 5μm.

